# Transcription-dependent DNA methylation at the imprinted *Zrsr1*-DMR

**DOI:** 10.1101/249524

**Authors:** Keiichiro Joh

## Abstract

*Zrsr1* is a paternally expressed imprinted gene located in the first intron of *Commd1*, and the *Zrsr1* promoter resides in a differentially methylated region (DMR) that is maternally methylated in the oocyte. However, a mechanism for the establishment of the methylation has remained obscure. *Commd1* is transcribed in the opposite direction to *Zrsr1* with predominant maternal expression, especially in the adult brain. In this study, we found *Commed1* transcribed through the DMR in the growing oocyte. *Zrsr1*-DMR methylation was abolished by the prevention of *Commd1* transcription. This result indicated that methylation at the *Zrsr1*-DMR was transcription-dependent.

## Introduction

Genomic imprinting is an epigenetic phenomenon of parent-of-origin–dependent expression that is observed in a subset of mammalian genes. Imprinted genes are expressed exclusively or predominantly from one of the two parental alleles, and are frequently located in clusters known as imprinted domains. The expression of genes in an imprinted domain is regulated by a discrete element called an imprinting center (IC) or an imprinting control region (ICR). Imprinted genes or imprinted domains have been found to link to differentially methylated regions (DMRs) that exhibit parent-of-origin–specific DNA methylation. Two classes of DMRs have been identified as follows: germline DMRs (gDMRs), or primary DMRs; and somatic DMRs (sDMRs), or secondary DMRs. gDMR methylation is established during gametogenesis, and sDMRs acquire methylation after fertilization under the direction of gDMRs. The ICs of the imprinted genes are located in their corresponding gDMRs. More than 20 gDMRs have been identified in mice, of which only three are paternally methylated (Arnaud, 2010, Barlow & Bartolomei, 2014, Ferguson-Smith, 2011, Kobayashi, Sakurai et al., 2012). Recent studies have identified an additional 11 new putative maternally methylated gDMRs (Wang, Zhang et al., 2014).

DNA methylation at gDMRs is the primary determinant of the allelic expression of imprinted genes, and the mechanisms of methylation establishment have been extensively investigated. The specific recruitment of *de novo* methylation machineries to gDMR methylation sites via the recognition of sequence elements and/or chromatin structures has been considered as a potential mechanism of germline-specific gDMR methylation establishment (Arnaud, 2010, Bartolomei & Ferguson-Smith, 2011). However, efforts to identify sequence motifs for gDMR methylation have not been successful. Several trans-acting factors for maternally methylated gDMRs have been found to be essential for the establishment of germline methylation in mice. For example, Dnmt3a has been identified as the enzyme responsible for *de novo* methylation of many maternal gDMRs (Hata, Okano et al., 2002, Kaneda, Okano et al., 2004). Dnmt3l, a DNA methyltransferase (DNMT)-like protein without enzymatic activity, is the likely co-factor of DNMTs (Bourc'his, Xu et al., 2001). Ablation of Kdm 1b, a histone demethylase of H3K4 di-and trimethylation, in oocytes resulted in the failure of methylation establishment at some maternal gDMRs (Ciccone, Su et al., 2009). In addition, the deletion of *Hira*, which encodes a histone H3.3 chaperon (Hira), led to global hypomethylation in oocytes (Nashun, Hill et al., 2015).

Kelsey et al. proposed a model for the establishment of methylation at the maternal gDMRs in the oocyte (Kelsey & Feil, 2013, Veselovska, Smallwood et al., 2015), which suggests that maternal methylation of gDMRs is regulated by the same mechanisms of general gene-body methylation reported for active genes (Ball, Li et al., 2009, Maunakea, Nagarajan et al., 2010). This was based on the findings that most maternal gDMRs are located in actively transcribed regions (Chotalia, Smallwood et al., 2009), that transcription is a prerequisite for the establishment of methylation at four maternal gDMRs (Chotalia et al., 2009, Singh, Sribenja et al., 2017, Smith, Futtner et al., 2011, Veselovska et al., 2015), and regarding the characteristics of the methylome and transcriptome in growing oocytes (Smallwood, Tomizawa et al., 2011, Veselovska et al., 2015). Analyses of the methylome revealed that most methylation in the oocyte genome occurs within actively transcribed regions, and that maternal gDMRs are not specifically targeted for methylation, but are instead methylated along with other parts of the transcribed regions where the gDMRs reside. Unlike the rest of the transcribed regions, gDMRs are likely protected against global demethylation during early embryonic development; only the gDMRs escape global demethylation and remain methylated for lifetime.

Methylation failed at the gDMRs in the *Gnas* locus and the KvDMR in the *Kcnq 1* locus when a poly(A) signal-truncation cassette was inserted into these loci to prevent transcription from elongation through the gDMRs (Chotalia et al., 2009, Singh et al., 2017). Failure of methylation was also reported at PWS-IC and the *Zac1-DMR* when the promoter regions from which transcription originated and then proceeded through the maternal gDMRs were deleted (Smith et al., 2011, Veselovska et al., 2015). Furthermore, most of these maternal gDMRs were located within transcribed regions in the growing oocyte (Chotalia et al., 2009). On the other hand, the establishment of sDMR- and lineage-specific methylation during the post-implantation stage clearly indicates the *de novo* methylation potency of early embryonic cells, including naïve/primed pluripotent stem cells (Monk, 2015). It is unknown whether *de novo* methylation occurs at gDMRs during the post-implantation stage after failure to establish gDMR methylation in the oocyte.

*Zrsr1* (*U2af1-rs1*) and *Commd 1 (Murr1)* are imprinted genes located in mouse proximal chromosome 11. *Zrsr1* is expressed ubiquitously in all adult tissues examined, and is expressed exclusively from the paternal allele. *Zrsr1* resides in the first intron of the *Commd1* gene and is transcribed in the opposite direction to the host gene. The *Zrsr1* promoter is located in a maternally methylated gDMR, i.e., the Zrsr1-DMR (Nabetani, Hatada et al., 1997). It is likely that maternal methylation at the Zrsr1-DMR causes imprinted expression by repressing maternal expression of the gene. *Commd1* is likewise expressed ubiquitously in adult mice, but is expressed from both parental alleles. However, *Commd1* expression from the maternal allele is stronger than expression from the paternal allele (i.e., predominant maternal expression), as exemplified in the adult brain (Wang et al., 2004). Although *Zrsr1-DMR* resides in the transcribed region of *Commd1*, the link between transcription and the establishment of methylation at this DMR has not been clarified.

## Results and Discussion

### Truncation of *Commd1* transcription results in methylation failure at Zrsr1-DMR in the growing oocyte

Methylation of maternal gDMRs of imprinted genes is established asynchronously during postnatal oocyte growth, typically between 5 and 25 days postpartum (dpp) (Lucifero, Mann et al., 2004). To verify that *de novo* methylation at Zrsr1-DMR is dependent on transcription resulting from *Commd1* expression in the oocyte, we analyzed *Commd1* expression and the methylation status of Zrsr1-DMR in growing oocytes from this period and in ovulated MII oocytes. *Commd 1* was expressed in all periods of oocyte maturation analyzed (Fig 1B). *De novo* methylation at Zrsr1-DMR started after 10 dpp and was completed between 15 dpp and maturation (Fig 1C). Thus Zrsr1-DMR was transcribed before and during the establishment of methylation. *Zrsr1* expression was not detected by RT-PCR during oocyte maturation (Fig EV1).

**Figure 1.**
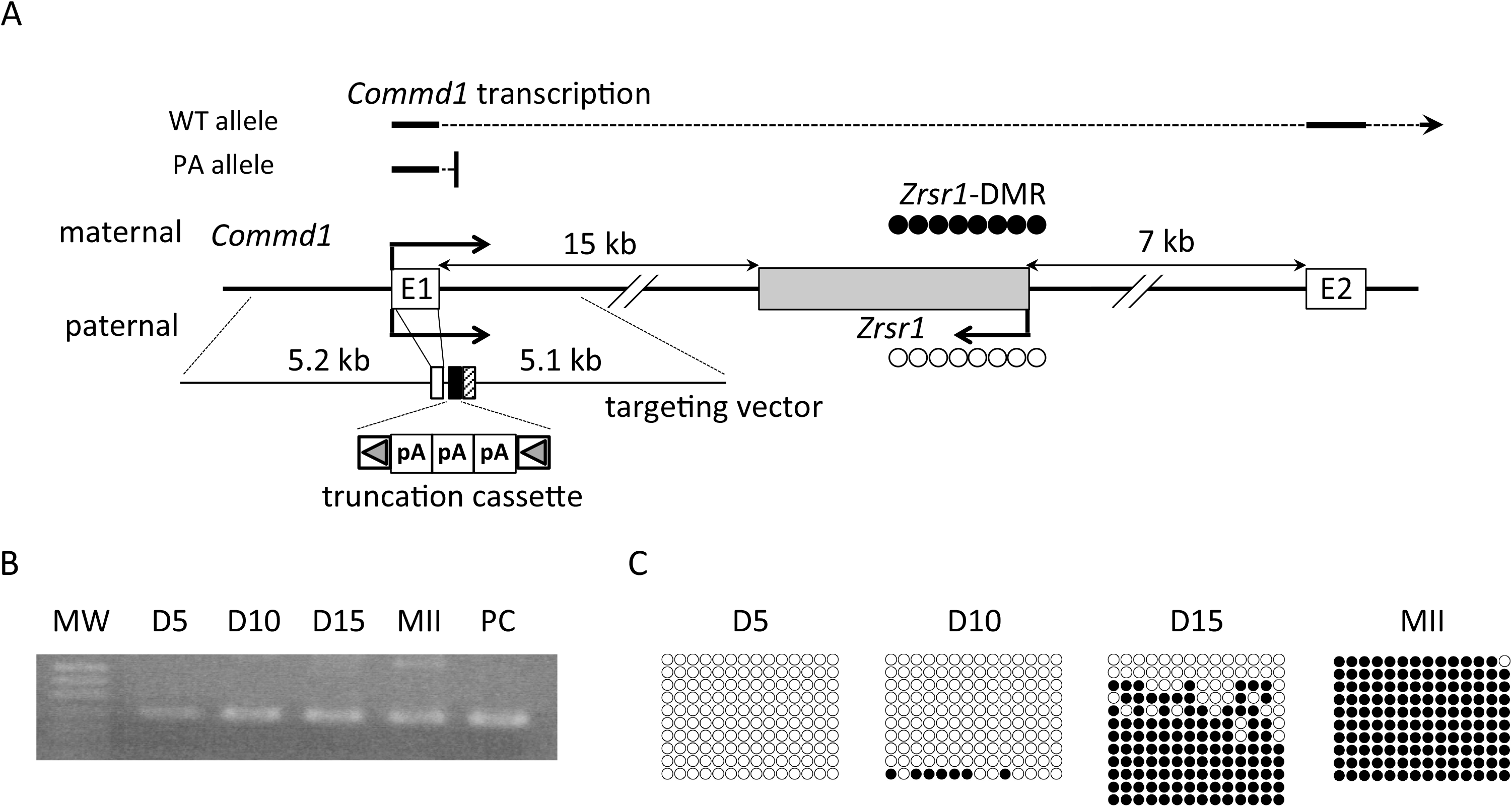
Structure of the *Zrsr1/Commd1* locus and analysis of *Commd1* expression and *Zrsr1*-DMR methylation in the oocyte. A *Zrsr1*, an approximately 2.8-kb intronless gene, and the first two exons of *Commd1* are represented by gray and white boxes, respectively. The schematic is not drawn to scale. Arrows above (maternal allele) and below (paternal allele) exon 1 and the *Zrsr1* gene represent the direction of transcription and the allelic expression status of the genes. The open and closed circles at the *Zrsr1* promoter indicate unmethylation and methylation, respectively. A schematic of the targeting vector is shown under the gene. The closed and hatched boxes represent the truncation cassette and the neo-selection marker gene, respectively. These elements are flanked by the 5.2-kb left arm containing exon 1, and the 5.1-kb right arm, which contains part of intron 1. The truncation cassette is flanked by loxP sites, represented by gray arrowheads enclosed in open rectangles. Expected transcription patterns of the WT and PA alleles are shown above the gene schematic. B RT-PCR analysis of *Commd1* expression in growing oocytes prepared from B6 female neonates at Day 5 (D5), Day 10 (D10), and Day 15 (D15) postpartum, and fully grown MII oocytes (MII) from B6 adult females. PC: positive control for RT-PCR using adult brain cDNA. MW: molecular weight marker. C Analysis of methylation at Zrsr1-DMR in growing and fully-grown oocytes used in (B). The 223-bp region in the DMR containing 14 CpGs was analyzed via bisulfite sequencing. Each row represents a dataset from one clone, and each circle represents one CpG site. Closed and open circles depict methylated and unmethylated CpGs, respectively.

To confirm that *de novo* methylation at Zrsr1-DMR was a prerequisite for *Commd1* expression, we inserted a truncation cassette containing three tandem copies of SV40 poly(A) signal into intron 1 of *Commd1* to generate the transcription-truncation allele *Commd1^PA^* (Fig 1A), and obtained *Commd1^+/PA^*-heterozygous mice in a C57BL/6J background. No *Commd1^PA/PA^* mice were born from the intercrossing of heterozygous parents. *Commd1^−/−^* mice have been shown to be embryonically lethal at E9.5 to E10.5 (van de Sluis, Muller et al., 2007); the absence of homozygous pups for the truncation allele was thus likely attributable to embryonic lethality, which strongly suggests that a truncation occurred as expected and rendered the *Commd1*^PA^ allele functionally null.

To assess the truncation of the *Commd1*^PA^ allele and *Zrsr1*-DMR methylation in the MII oocyte, MII oocytes were obtained from adult F1 females generated from the cross between *Commd1^+/PA^* B6 females and WT PWK males. *Commd1^PA(B6)/+(PWK)^* mice, termed PA F1 mice, and *Commd1^+(B)/+(PWK)^* mice, termed WT F1 mice, were obtained from F1 littermates. The allelic expression of *Commd1* was analyzed by RFLP analysis of RT-PCR products with the primers Comm-F1 and Comm-R1, located at exon 1 and exon 2, respectively. Expression of the *Commd1^PA^* allele was not detected in MII oocytes prepared from the PA F1 females, although expression of the PWK allele was detected. In contrast, both alleles were expressed in oocytes from WT F1 female littermates (Fig 2A). The Zrsr1-DMR was completely unmethylated on the truncated allele (B6) in the MII oocytes from the PA F1 females, in contrast to the WT PWK allele, which was completely methylated. As expected, in oocytes from the WT F1 females, Zrsr1-DMR was completely methylated on both the B6 and PWK alleles (Fig 2B). These results indicate that transcription termination occurred in intron 1, likely at the truncation cassette, and resulted in the loss of transcription through the DMR, which led to methylation failure at the DMR during oogenesis. The results indicate that transcription through the *Zrsr1-DMR* is essential for the establishment of methylation at the DMR in the growing oocyte, as reported for four other imprinted loci (Chotalia et al., 2009, Singh et al., 2017, Smith et al., 2011, Veselovska et al., 2015).

**Figure 2.**
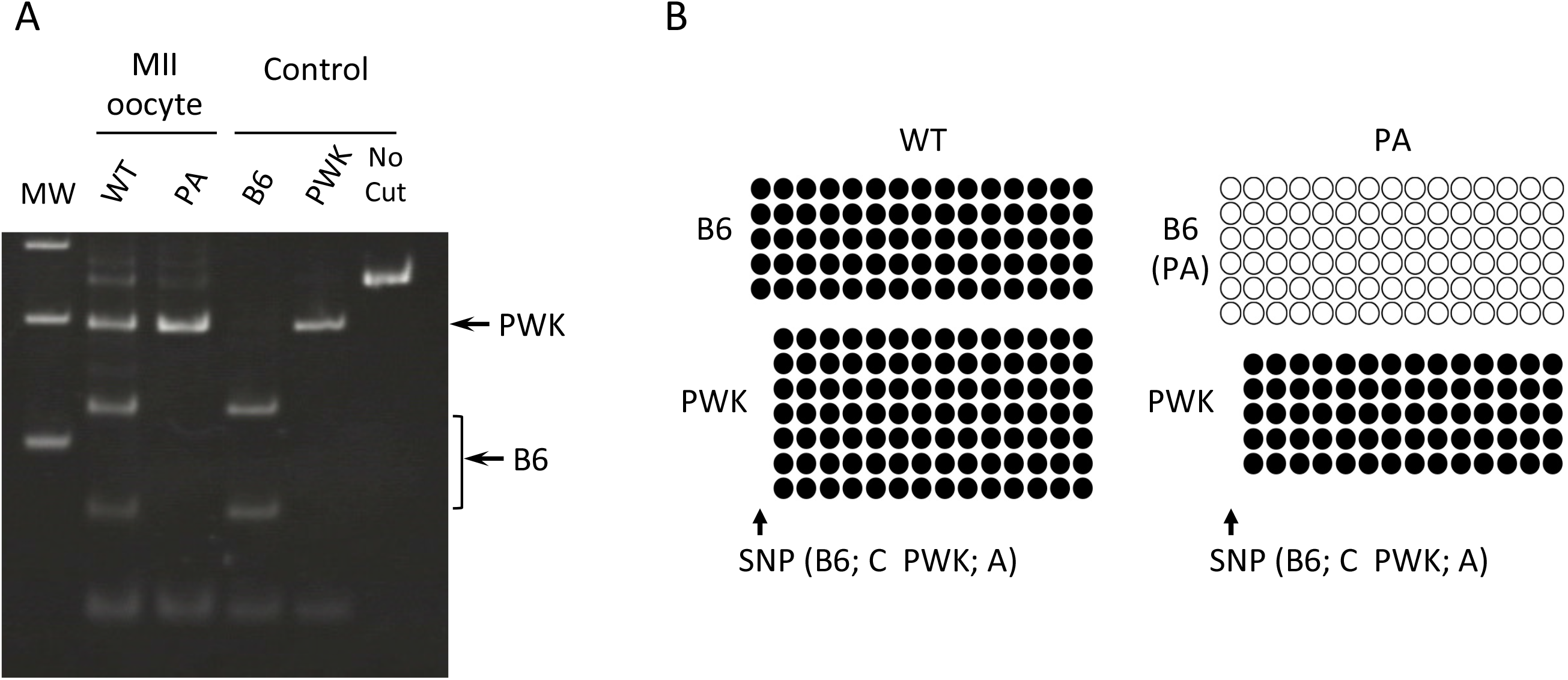
Analysis of allelic *Commd1* expression and *Zrsr1*-DMR methylation in PA and WT MII oocytes. A MII oocytes were prepared from *Commd1^PA(B6)/+(PWK)^* females (PA) and *Commd1^+(B6)/+(PWK)^* females (WT). RT-PCR was performed as in Figure 1B. Amplified cDNA (250 bp) was digested with NlaIII (CATG). There are two NlaIII restriction sites in the B6 amplicon, but one of these is absent in the PWK amplicon because of a single nucleotide polymorphism (SNP) between the two strains. Adult brain RNA from each of the strains was used as the control. MW: molecular weight marker. B A 278-bp region in Zrsr1-DMR containing 15 CpG sites (B6) or 14 CpG sites (PWK) was analyzed via bisulfite sequencing. The alleles were discriminated between via an SNP (C in B6 and A in PWK) indicated as the leftmost CpG site in the B6 sequence.

## Materials and Methods

### Generation of *Commd1-PA* mice

A truncation cassette was constructed by cloning three copies of the SV40 poly(A) signal from the expression vector pGFP-N1 (Clontech) into pT7Blue (Novagen). The truncation cassette was inserted at the genomic site 23 bp downstream of exon 1 using the gene-targeting method (Fig 1A). The targeting construct was generated by homologous recombination with a truncation cassette clone and a mouse *Commd1* BAC clone (BAC RP24-216A32, BACPAC Resources) in *E. coli* as previously described (Zhang, Muyrers et al., 2000). The targeting construct contained the following mouse *Commd1* genomic sequences: a 5.2-kb 5' sequence containing exon 1, and a 5.1-kb 3'-sequence containing a part of intron 1. An embryonic stem cell (ES) clone with the truncation cassette inserted in the precise genomic position was identified by Southern blotting and PCR analyses of genomic structure, and was used to generate chimeric *Commd1-PA* mice. The *neo* gene in the targeting vector was flanked by FRTs and removed from the *Commd1*-truncated mice via Flippase expression. *Commd1-PA* mice in a C57BL/6J (denoted B6) genetic background was obtained by backcrossing with wild type (WT) C57BL/6J mice for five generations. This study was approved by the Ethical Committee for Animal Experiment of Saga University.

### Preparation of primordial germ cells, metaphase II oocytes, and blastocysts

Mouse female primordial germ cells (PGC) were prepared from Day 5, Day 10, and Day 15 female C57BL/6 neonates as previously described (Kobayashi, Sakurai et al., 2013). The metaphase II (MII) oocytes were obtained from superovulated six- to eight-week-old female mice using the procedure described by Nakagata *et al.* (Nakagata, Takeo et al., 2013). Blastocysts were prepared by *in vitro* fertilization (IVF) using MII oocytes from *Commd1^PA+^* B6 mice and sperm from 12- to 14-week-old CAG-Cre transgenic BALB/c mice. After cumulus-oocyte complexes had been coincubated with sperm for 3 h, fertilized oocytes were cultured to the blastocyst stage at 37 °C and 5% CO_2_ in humidified air for 96–120 h, as previously described (24).

### DNA preparation and methylation analysis

Approximately 200 growing oocytes and MII oocytes were lysed in lysis buffer (0.5% SDS, 250 ng/"l proteinase K, 100 ng/μl yeast tRNA) at 37 °C for 60 min, and DNA in the lysate was bisulfite-converted by treating the lysate with the EpiTect Bisulfite Kit (QIAGEN, #59104). Recovered DNA was used for amplification of Zrsr1-DMR via nested PCR. Amplified DNA was cloned in pT7Blue T-vector (Novagen, #69820) and the resulting sequences were analyzed with BigDye Terminator v3.1 (Applied Biosystems, #4337455) on an ABI3130 sequencer (Applied Biosystems). Blastocyst DNA was prepared from the precipitate of ISOGEN II lysate during RNA preparation with ISOGENOME (NIPPON GENE, #318-08111). DNA from adult tissues was prepared using QIAamp DNA Mini Kit (QIAGEN, #51306). Blastocyst DNA was genotyped by genomic PCR, using primers on both sides of the truncation cassette (Fig S2). DNA from tissues was also bisulfite-converted using the EZ DNA Methylation Kit (Zymo Research Corp., #D5001). PCR, cloning, and sequencing were performed as described for oocyte DNA.

### RNA preparation

RNA was prepared from approximately 100 growing and MII oocytes using the RNeasy Micro Kit (QIAGEN, #74004). For RNA preparation from blastocysts and adult tissues, samples were lysed with ISOGEN II (NIPPON GENE, #311-07361), and the cleared lysates were recovered by centrifugation according to the manufacturer's instructions. Three blastocysts were pooled for RNA preparation, and the lysate was loaded onto a spin column from the RNeasy Mini Kit (QIAGEN, #74104). DNase I treatment, column wash, and RNA elution were performed according to the manufacturer's instructions.

### Expression analysis

To analyze *Commd1* expression in oocyte and adult tissues, cDNA was synthesized with random primers (Takara, #3802) and reverse transcriptase (TOYOBO, TRT-101), and the *Commd1* cDNA was amplified by PCR with the primers Comm-F1 (exon 1) and Comm-R1 (exon 2). To analyze allelic expression, restriction fragment length polymorphism (RFLP) analysis was performed via NlaIII digestion of the amplified cDNA. Allelic expression of *Zrsr1* was analyzed by BigDye terminator sequencing of the cDNA amplified with the primers Zrsr-F1 and Zrsr-R1. No rs numbers were assigned to the single nucleotide polymorphisms (SNPs) that were used to discriminate between the C57BL/6J and PWK alleles.

## Acknowledgements

We thank Dr Tatsuya Kishino from the Gene Research Center, Center for Frontier Life Sciences, Nagasaki University, for providing the PWK mice used in this study.

## Figure legends

**Figure S1.**
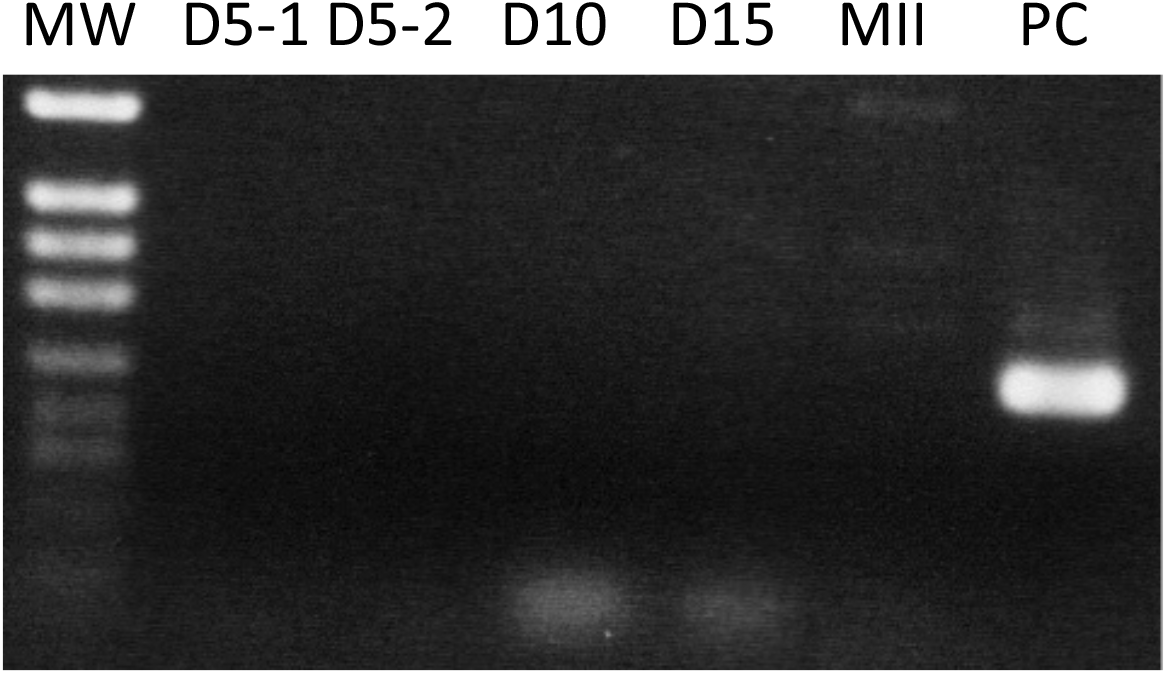
RT-PCR analysis of *Zrsr1* expression in growing oocytes. RT-PCR was done with primers Zrsr-F1 and Zrsr-R1 using cDNAs in Fig 1B. Growing oocytes were prepared from B6 female neonates at Day 5 (D5), Day 10 (D10), and Day 15 (D15) postpartum, and fully grown MII oocytes (MII) from B6 adult females. PC: positive control for RT-PCR using adult brain cDNA. MW: molecular weight marker. Two different cDNA batches were used for D5 RNA.

